# A FINITE ELEMENT FRAMEWORK FOR BULK-SURFACE COUPLED PDES TO SOLVE MOVING BOUNDARY PROBLEMS IN BIOPHYSICS^*^

**DOI:** 10.1101/2025.10.27.684936

**Authors:** Alessandro Contri, André Massing, Padmini Rangamani

## Abstract

We consider moving boundary problems for biophysics and introduce a new computational framework to handle the complexity of the bulk-surface PDEs. In our framework, interpretability is maintained by adapting the fast, generalizable and accurate structure preservation scheme in [29]. We show that mesh distortion is mitigated by adopting the pioneering work of [36], which is tied to an Arbitrary Lagrangian Eulerian (ALE) framework. We test our algorithms accuracy on moving surfaces with boundary for the following PDEs: advection-diffusion-reaction equations, phase-field models of Cahn-Hilliard type, and Helfrich energy gradient flows. We performed convergence studies for all the schemes introduced to demonstrate accuracy. We use a staggered approach to achieve coupling and further verify the convergence of this coupling using numerical experiments. Finally, we demonstrate broad applicability of our work by simulating state-of-the-art tests of biophysical models that involve membrane deformation.

## 1. Introduction

Many biophysical phenomena in cell biology can be characterized by the feedback between form and function. Cells routinely change their shape during cell motility, wound healing, and development. Such shape changes are a result of coordination of multiple biochemical and mechanical processes within the cell. Not surprisingly, these phenomena have been studied extensively using models based on partial differential equations (PDEs) with different levels of biophysical complexity [78]. In concert with experiments, such models have provided insight into the mechanisms underlying the change in cell shape [31, 95, 63]. However, an open challenge in the field of computational biophysics is the development of robust computational tools for numerical solutions of the governing PDEs. For example, cellular membranes are mechanical boundaries that are characterized by lateral dimensions that are large compared to their thickness, appearing in a variety of structures in cells [99]. It has been proven that the properties of these surfaces are crucial for the genesis and maintenance of cellular membrane systems [16, 86]. These restructuring phenomena strongly depend on the components of the membranes themselves and their surroundings [19, 10, 96, 17, 73, 104, 2, 5, 98], leading to PDE systems coupling complex multiphysics to free surfaces problems. The resulting systems typically exhibit highly non-linear dynamics and a general numerical framework to simulate these rich dynamics in a unified and stable setting is still missing. The goal of this article is to present a finite element based computational framework which addresses some of the key challenges arising in continuum-mechanics based computational models of cells.

### 1.1. State of the art

Numerous numerical methods have been proposed to deal with the challenges posed by a moving deformable membrane [61, 92, 1, 71, 93, 69, 90, 85, 7], including: immersed boundary methods, level-set method, mesh-free methods, particle methods, and the phase-field method. Despite numerous successes, even the most advanced schemes seem to lack the following features:

P.1 It is not rare, in the targeted applications, to incur advection-dominant (AD) equations possibly posed on a *moving surface*. Characteristic examples can be the transport of surfactants in two-phase flow or the interaction between actin’s barbed ends and the cellular membrane. There exists a large bulk of literature for parabolic advection-diffusion-reaction (ADR) PDEs on moving surfaces [38, 40, 18, 80, 39, 67, 68, 42, 25]. For what concerns AD equations, the bulk setting has been extensively explored and the surface case has seen recent developments [91, 22, 97, 23, 33, 100, 101]. Surprisingly, most afore-mentioned approaches only consider the case of stationary surfaces, leaving the AD case largely unaddressed. A notable exception is the work in [53], where characteristic cut finite element methods on moving surfaces is proposed.
P.2 Physical structure preservation (SP) is still overlooked by the majority of the available articles. It is crucial, in order to maintain physical interpretability as well as numerical stability, to be able to guarantee constraints such as: mass and density positivity, phase bounds, energy decay, mass preservation, etc. A vast and diversified literature exist dedicated to preserve bounds, see [79, 74, 34, 75, 103, 102, 76, 28, 58, 56] and references therein. Unfortunately, this richness of options is lacking if we consider PDEs on surfaces, let alone moving surfaces.
P.3 Most articles assume idealized geometries, in which the complexity is still highly reduced with respect to realistic geometries resulting from imaging techniques [72]. This is especially crucial in this context since we expect the domain evolution to be dependent on the coupled equations and not known *a priori*. Steps are continuously being taken in this direction for what concerns geometric flows, which notoriously drive meshes towards bad conditioning, see [11, 81, 57, 36, 9, 51]. However, even in these cases, one is quite far from the structures considered in more applied articles such as [50].

The present work aims to propose a first step towards a cure of the missing pieces described above. It is meant to bridge the complexity gap between numerical accuracy and biophysical complexity in moving domains scenario. While there exist a plethora of numerical methods which each target a specific problem class or particular issue, we here employ and extend some of the most recent state-of-the-art methods and combine them into a holistic simulation framework which can handle coupled multiphysics and shape deformation problems arising in computational cell biology.

### 1.2. New contributions and outline

In Section 2, we provide the notation necessary for the formulation and discretization of PDEs posed on evolving domains. For purposes of generality, we keep the notation the same in the bulk and on the surface, only distinguishing between the two as needed. In Section 3, we adapt the structure-preserving scheme in [30] to moving bulk and surface problems as a solution to P.2. The model-agnostic, point-wise approach of [30] is fit to the first-order, piecewise-linear finite element framework here used. The result is a fast, generalizable and flexible scheme that can be readily adapted to multiple PDEs posed on moving bulk and surfaces. In this context, we use it to bound the PDE solution to physically-relevant values while at the same time preserving the total mass. In Section 4, we introduce a mesh redistribution algorithm based on the surface tangential motion of [36] and an Arbitrary Lagrangian Eulerian (ALE) approach. The two-step scheme in [36] is decoupled from the gradient flow dynamics and applied to general motions. This allows us to greatly alleviate the need of remeshing when simulating domains characterized by large deformations. At the same time, the scheme does not interfere with the shape evolution, maintaining accuracy. Both schemes in Section 3 and Section 4 take the form of a post-processing step and thus have the advantage of being a tunable feature of the code. In Section 5 we augment an ADR scheme for moving surfaces with a continuous interior penalty (CIP) stabilization to handle AD cases and solve P.1. The stabilization is proven to be convergent and so is its coupling with the structure-preserving algorithm of Section 3. The flexibility of the setting is further verified with an additional real-case example. In Section 6 we apply structure preservation to a Cahn-Hilliard type equation on moving surfaces. Numerical tests are presented that highlight how the algorithm in Section 3 is readily applicable to moving surfaces and true to the underlying physics. In Section 7, we focus on P.3, applying the mesh redistribution scheme of Section 4 to the parametric method of Barrett, Garke and Nürnberg [14] for Helfrich flow. The postprocessing step is extensively tested with state-of-the-art examples revealing good convergence properties and long-term stability. In Section 8, we couple the solvers in Section 5 through 7 together in a staggered approach. Convergence is tested with manufactured solutions [67] where the ADR algorithm and a simplified version of the Helfrich algorithm are coupled. In the same context a PDE system modeling tumor growth is also reproduced. The applicability of our approach to relevant cell biophysics is further explored simulating phase separation on a deformable membrane, where the Cahn-Hilliard and Helfrich schemes are coupled.

## 2. Model

This article explores numerical approaches to biophysical cell phenomena where the dynamics involve moving boundaries, such as cell membranes, that are assumed to have negligible thickness and thus are treated as moving hypersurfaces. As a result, we begin by outlining the essential notation required for both formulating and discretizing partial differential equations on evolving domains, with specific attention given to surface-bound PDEs [39, 15].

Let *M* ⊂ ℝ^*d*^, *d* = 2, 3, be an *m*-dimensional oriented, compact, *C*^2^ manifold with boundary *∂M*. The notation Ω ≡ *M* when dim (*M*) = *d* and Γ ≡ *M* when dim (*M*) = *d*− 1 will also be used. In the case dim(*M*) = *d*− 1, we denote the tangent space with respect to ***p*** ∈ *M* with *T*_***p***_(*M*) and the associated tangent bundle with *TM*. The *unit normal vector* ***n***_*M*_ (***p***) to *M* at ***p*** is defined as the vector ***n***_*M*_ (***p***) ∈ ℝ^*d*^ such that ***n***_*M*_ (***p***) ⊥ *T*_***p***_(*M*), ∥***n***_*M*_ (***p***)∥ = 1 and ***n***_*M*_ (***p***) agrees with the orientation given. The norm ∥·∥ is the standard Euclidean norm in ℝ^*d*^. For dim (*M*) = *d* − 1 we define the *tangential projection* at ***p*** as

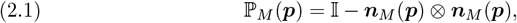

where 𝕀 is the identity matrix. For dim (*M*) = *d*, we set ℙ_*M*_ (***p***) = 𝕀 and ***n***_*M*_ (***p***) = **0**. This allows us to define the *tangential gradient* of a differentiable function *f* : *M* → ℝ at ***p*** as

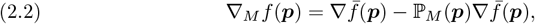

where 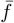 is a smooth extension of *f* to a *d*-dimensional neighborhood of *M* such that 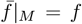 and ∇ is the Euclidean gradient in ℝ ^*d*^. Analogously, the *tangential divergence* is defined as ∇_*M*_ · *f* (***p***) = Tr(∇_*M*_ *f* (***p***)). This leads to the definition of the Laplace-Beltrami operator for a *C*^2^-function *f* : *M* → ℝ as

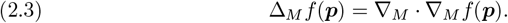

For ***f*** : *M* → ℝ^*d*^ and 𝔽 : *M* → ℝ^*d×d*^ we define (∇_*M*_ ***f***)_*ij*_ = (∇(***f*** ·***e***_*i*_))_*j*_ and (∇_*M*_ ·𝔽)_*i*_ = ∇_*M*_ · (𝔽^*T*^ ***e***_*i*_) = Tr(∇_*M*_ (𝔽^*T*^ ***e***_*i*_)), where ***e***_*i*_, *i* ∈ *{*1, …, *d}* is the canonical basis in ℝ^*d*^. This allows us to define Δ_*M*_ ***f*** = ∇_*M*_ · ∇_*M*_ ***f***. In order to analyze curvature properties of *M*, we introduce the *extended Weingarten map* ℍ as

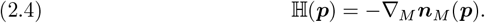

It can be shown that ℍ is symmetric, has an eigenvalue 0 in the direction of the normal, and restricts to the Weingarten map 𝕎(***p***) on the tangent space *T*_***p***_*M*. For ***p*** ∈ *M*, the *mean curvature* is defined as

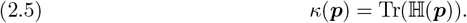

We recall that all of the above definitions are independent of the extension 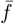. A sketch of possible model setups is shown in Figure 1.

**Fig. 1:**
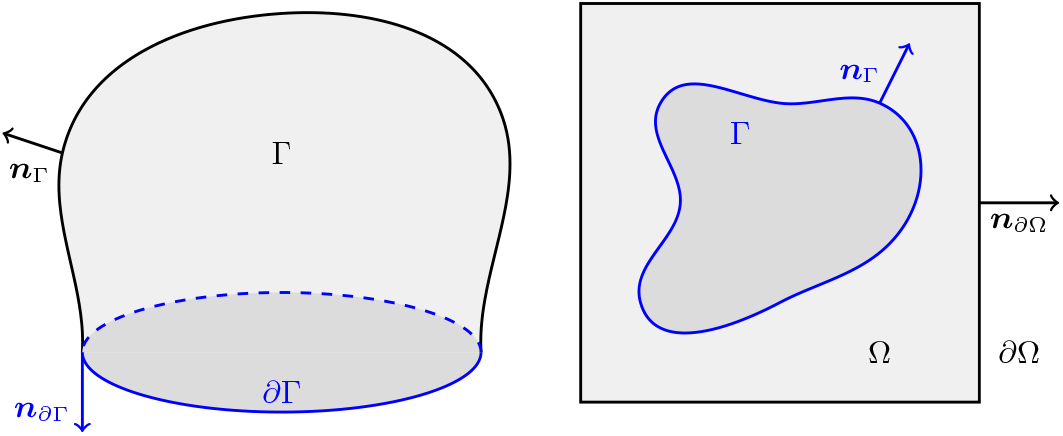
Sketch of an open and closed geometry for notation purposes. One is a 3D object and the other is a 2D object.

We consider a time interval *J* = [0, *T*], *T >* 0 and a *C*^2^-*evolving manifold {M* (*t*)*}*_*t*∈*J*_ in ℝ^*d*^. In this article an evolving manifold is modeled by a *reference manifold* 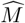 together with a *flow map*

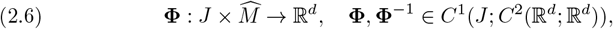

such that

- denoting 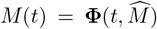, the map 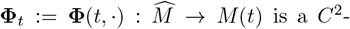 diffeomorphism with inverse map 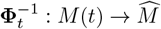,
- 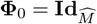, where 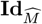 is the identity map on the reference manifold.

The mapping **Φ**_*t*_ can be used to define pull-back and push-forward maps of functions [4, 3]

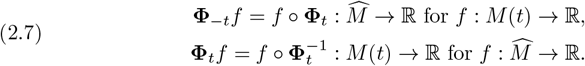

Often we will call the space-time set 𝒢_*T*_ = ∪_*t*∈*J*_ (*{t} × M* (*t*)) a *C*^2^-*evolving manifold* and identify it with *{M* (*t*)*}*_*t*∈*J*_. In addition, we assume that there exists a velocity field ***v*** : *J ×* ℝ^*d*^ → ℝ^*d*^ with ***v*** ∈ *C*^0^(*J*; *C*^2^(ℝ^*d*^; ℝ^*d*^)) such that for any *t* ∈ *J* and every 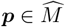

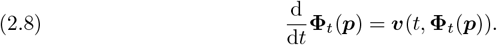

For ***x*** ∈ *M* (*t*), the velocity ***v*** can be split into a tangential component ***v***^⊤^(*t*, ***x***) ∈ *T*_***x***_*M* (*t*) and a normal one ***v***^⊥^(*t*, ***x***) = ***v***(*t*, ***x***) − ***v***^⊤^(*t*, ***x***).

This allows us to define a *normal derivative* of a scalar function *f* on *M* (*t*) as

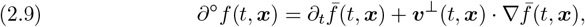

where 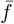is a smooth extension of *f* in a space-time neighborhood of *M* (*t*). The *material derivative* is defined as

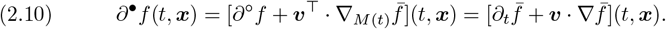

The definition of a flow map is not unique and the same evolving manifold can be described by different maps. The computational domain’s flow map can be chosen so to maintain good mesh properties while keeping the original map **Φ** for the PDE evolution. This is achieved by introducing a second flow map **Φ**^𝒜^ called *Arbitrary Lagrangian Eulerian (ALE) map* with corresponding velocity ***v***^𝒜^ and ALE material derivative *∂*^𝒜^. It can be shown that ***v***^𝒜^ (*t*, ***x***) − ***v***(*t*, ***x***) ∈ *T*_***x***_*M* (*t*) and that

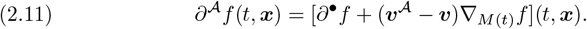

It is worth noting that the normal velocity ***v***^⊥^ is in any case uniquely determined, i.e. ***v***^⊥^(*t*, ·) = ***v***^𝒜,⊥^(*t*, ·), *∀t* ∈ *J*, including the normal velocity of the boundary *∂M* (*t*).

### 2.1. Space discretization

The geometries are discretized using piecewise linear elements, where for surfaces we follow the setup of [39]. An *m*-dimensional manifold *M* is approximated by a triangulated *m*-dimensional domain denoted by *M*_*h*_. The elements of the discretization will be distinguished based on their codimension. We suppose *M*_*h*_ is composed by a collection 𝒯_*h*_ of *m*-simplices which vertices ***y***_*i*_, *i* = 1, …, *N* lie on *M* and such that

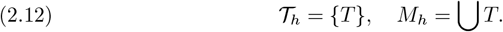

For each *T* ∈ 𝒯_*h*_ we denote by *h*(*T*) its diameter and define the *mesh-size* as

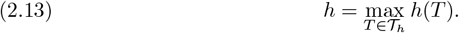

The boundary of every element *T* is composed by *m* + 1 facets of dimension *m* − 1, forming a collection ℱ = *{F}*. The discrete boundary of *M*_*h*_ is defined as the union of those facets which defining points lie on *∂M* and is denoted *∂M*_*h*_. It is useful to distinguish between *internal* and *boundary* facets, defined as

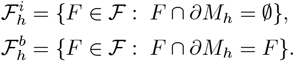

Every element *T* has a (constant) normal which we will denote by ***n***_*T*_. In the case dim(*M*_*h*_) ≡ *d* we define ***n***_*T*_ ≡ 0. In the case dim(*M*_*h*_) ≡ *d* − 1 we define a discrete projection ℙ_*T*_ = 𝕀 − ***n***_*T*_ ⊗ ***n***_*T*_ and relative tangential gradient, understood in an element-wise sense. The facet normal, which we denote ***n***_*F*_, is well-defined once we consider it on the boundary of an element *T*. For this reason for each *F* ∈ *∂T*, ***n***_*F*_ is the unit co-normal vector perpendicular to both ***n***_*T*_ and *F* and directed outward with respect to the element *T*. We will also use the notation 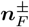, where given as *T* ^+^ and *T*^−^ the neighboring elements to *F*, then 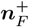 is the normal as seen from *T* ^+^ and 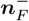 is the normal as seen from *T*^−^. In the bulk case we simply have that 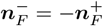, instead for surface cases the two normals are not co-planar. An illustration of these objects is given in Figure 2. We will again use the notation Ω_*h*_ and Γ_*h*_ when we need to distinguish between *d*-dimensional and (*d −* 1)-dimensional discrete manifolds, respectively.

**Fig. 2:**
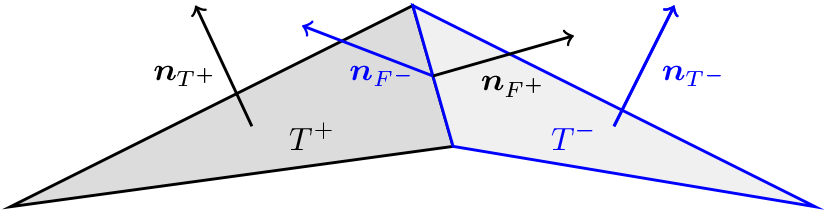
Sketch of an discrete discrete mesh entities of a discrete surface *M*_*h*_.

Throughout this work, we use a first-order fitted finite element method to discretize the PDEs in space [46]. In particular, for surfaces, we employ the surface finite element method (SFEM) as presented in [39]. The corresponding finite element space is defined as

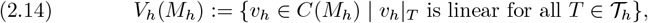

which can be described as the span of *N* piecewise linear continuous basis functions *ϕ*_*i*_, *i* = 1, …, *N* such that *ϕ*_*i*_(***y***_*j*_) = *δ*_*ij*_. Occasionally, we will also need the space

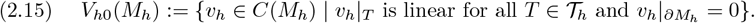

Integration over mesh entities follows the same notation as for the continuous one, where the summation over every element of the collection is implicit, i.e. we write

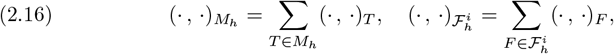

where (·,·) is a certain inner product. The same element-wise summation is used in case of discrete norms.

### 2.2. Time discretization

To discretize the evolving manifold 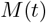 we use the reference discretization 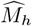 and transport it using a suitable ALE map **Φ**^𝒜^ [55, 60, 89, 52, 49, 8, 18, 68, 3]. The reference points 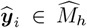 defining the simplices of the reference triangulation 𝒯_*h*_ are evolved as 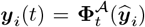. This allows to define evolved elements *T* (*t*) and facets *F* (*t*) such that

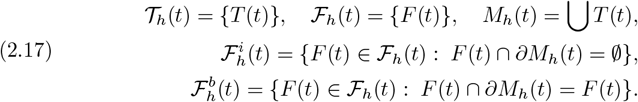

For the discretization of the function spaces, we start with the finite element space defined in (2.14), then transport the basis functions using the flow map 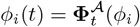 and define the evolved finite element space as their span

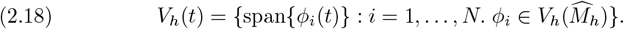

In an analogous way we can also define *V*_*h*0_(*t*). It follows from the definitions that 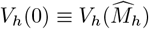. Discrete material velocities are defined by

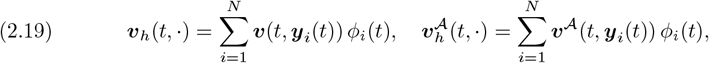

with corresponding derivatives

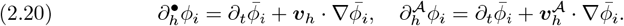

Using an ALE map, which describes the motion of the computational domain, we have that 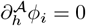.

In this work, time-dependent PDEs are discretized in a collection of time points *{t*_0_, …, *t*_*N*_*}*. We denote quantities on discrete time points by a superscript, where (·)^*n*^ refers to the previous time step and (·)^*n*+1^ refers to the current time step. The time step size is defined as *τ* ^*n*+1^ = *t*_*n*+1_ − *t*_*n*_ and for the sake of simplicity we restrict in what follows to constant time step size *τ*, with the understanding that every first-order-in-time scheme is amenable to adaptive time stepping.

#### Remark 2.1.

From now on, we will often omit the explicit dependence on time, indicating for example *M*_*h*_(*t*) as simply *M*_*h*_ and leaving it to the context to indicate the considered time.

## 3. Structure preservation

In this section we describe how to deal with the challenges highlighted in P.2. In the effort of maintaining the interpretability of the considered systems, we focus on bounds and mass preservation. The method recently proposed in [30, 29] provides a flexible approach to achieve such goals for a wide variety of finite element formulations on stationary bulk domains. Here, we extend those advances to moving surfaces and moving bulk domains. The main advantages of the employed method are:

- Bounds preservation is achieved in such a way that, in its simplest form, it reduces to the widely used cut-off approach.
- Accuracy is theoretically retained for higher order methods in space and time, granted some underlying hypotheses are satisfied. This is not the case, for example, for discrete maximum principle preserving schemes.
- The approach can easily be incorporated as a post-processing step without requiring explicit time integration. While explicit time integration as reviewed in [103] can be effective for hyperbolic equations, here we are also interested in parabolic equations.
- The simplicity of the algorithm makes it suitable for implementation in legacy codes with negligible overhead computational time with respect to the original non-preserving discretization method. Nevertheless, the method enjoys good stability properties and even allows for error analysis for some specific cases, see [29].
- No *ad hoc* problem reformulation is needed, but the scheme only acts on the nodal values of the solution through a Lagrange multiplier approach.

Following the presentation in [29], the derivation of the method starts from a nonlinear PDE in the form

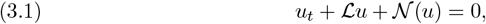

where ℒ is a linear or non-negative operator and 𝒩 (*u*) is a semilinear or quasi-linear operator. The operators ℒ and 𝒩 (*u*) are acting on functions defined on the space-time manifold 𝒢_*T*_. As underlying assumption, the solution to the continuum problem lies in the interval [*a, b*], i.e. given *a* ≤ *u*(0, ***p***) ≤ *b* for all 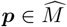 then *a* ≤ *u*(*t*, ***x***) ≤ *b* for all (*t*, ***x***) ∈ 𝒢_*T*_. The corresponding generic spatial discretization is

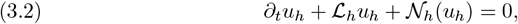

where we assume *u*_*h*_ ∈ *V*_*h*_. To make the scheme bound-preserving, a Lagrange multiplier *λ*_*h*_ is introduced together with the quadratic function *g*(*u*) = (*b− u*)(*u − a*). The problem is then reformulated as

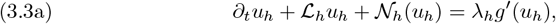

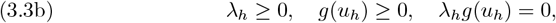

where the second line represents the usual Karush-Kuhn-Tucker conditions for constrained optimization [62, 70]. A core assumption of the scheme presented in [29] is that (3.3) is satisfied *pointwise* in a set of points Σ_*h*_ that can be both mesh points or collocation points for *M*_*h*_. We must then have:

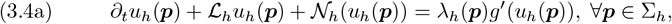

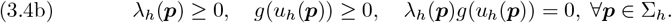

It is important to notice that Σ_*h*_ does not include points where essential boundary conditions are applied. The next step is to discretize (3.4) in time, where we here focus on the case of Backward-Euler (BE or BDF-1) time integration in the interval {*t*^*n*^, *t*^*n*+1^}. Higher order extensions are presented in [29] and can be adapted following similar arguments. The key idea is to apply an operator splitting approach to (3.3) dividing the problem in two steps.

In the *predictor* step we solve the unconstrained problem

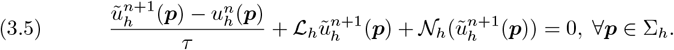

In the *corrector* step we solve the constrained problem

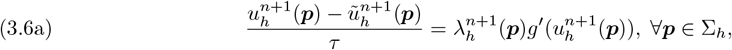

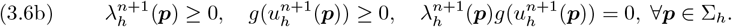

If we are only interested in bound preservation, it turns out that the solution to (3.6) is the cutoff function

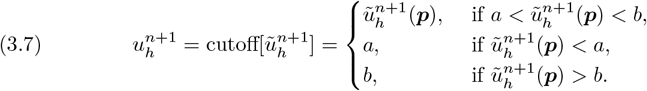

The above step is unfortunately not mass-preserving even if the solution is. To fix this, the authors in [30, 29] introduce an additional global space-independent Lagrange multiplier 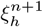. The corrector step (3.6) is modified to include the mass preservation constraint,

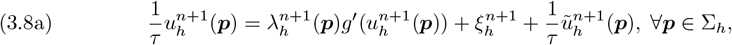

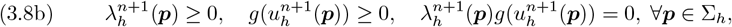

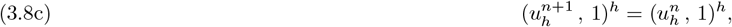

where (·, ·)^*h*^ is a suitably chosen discrete inner product. In our case this discrete inner product, that has to be expressed pointwise as 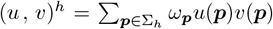 is assumed to be equivalent to the classical *L*^2^ inner product on *V*_*h*_. For this reason, whenever required, we will choose (·,·)^*h*^ to be the mass-lumped inner product. Rearranging (3.8a) as

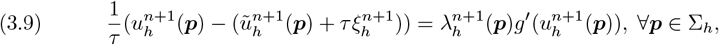

the solution is then 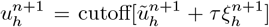 where 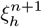 is the solution to the nonlinear equation

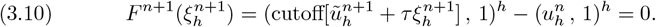

Bounds on the energy norm are provided for a class of non-linear PDEs in [29] together with error analysis of a restricted class of problems.

### Remark 3.1.

Since (*F* ^*n*+1^)^*′*^ might not exist, the authors of the scheme in [30, 29] suggest the use of a secant method to find the solution of *F* ^*n*+1^(*ξ*) = 0 in (3.10) using

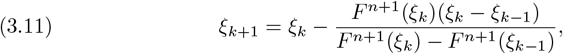

together with initial guesses *ξ*_0_ = 0 and *ξ*_1_ = 𝒪(*τ*). From a practical standpoint, the algorithm converged in less than four iterations for each timestep for the case at hand.

The technique introduced is clearly directly applicable to a wide variety of equations. The authors themselves in [30, 29] have proposed experiments for Allen-Cahn, Cahn-Hilliard with variable mobility and Fokker-Planck equations on a fixed bulk domain. Inspired by the biophysical mechanisms driving cell reshaping, we will demonstrate their accuracy when modeling complex advection-diffusion-reaction equations and phase-separation phenomena. The structure preservation properties will allow to simultaneously maintain stability, interpretability and physical complexity.

## 4. Tangential grid control and ALE discretization

When dealing with evolving domains in a fitted framework, it is not uncommon to incur deformations that lead to highly distorted meshes. This is especially true when gradient flows are involved, since they only prescribe the velocity and hence the displacement in normal direction. Following 2.11, we introduce a particular ALE map tailored for gradient flow dynamics that works as follows.

Consider the time interval *{t*_*n*_, *t*_*n*+1_*}*, and suppose the surface 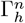 would be displaced, following the physics of the problem, to an updated surface 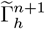 at time *t*_*n*+1_ which is highly distorted. We define an artificial tangential motion aimed at maintaining the surface shape (and consequently energy) but redistributing the nodes more favorably to reduce mesh distortions. We choose the two-stage algorithm proposed in [36] for the following reasons:

- The method has proven to be very effective for gradient flows such as the mean curvature flow and the surface diffusion flow.
- Its two-stage nature makes it a tunable feature of the code, that can be turned on and off at will.
- Being a two-stage process and not embedded in the gradient-flow algorithm, it can be applied on the mesh evolution independently of the presence of the gradient flow.
- Beyond the scope of the current work, the method can potentially be applied to higher order in time integration schemes.

The algorithm presented in [36] is based on the assumption that the starting mesh is regular enough. The idea is then to impose an artificial tangential velocity that requires 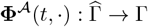 to be an harmonic map, i.e.

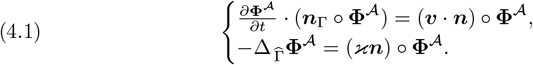

Equivalently, **Φ**^𝒜^ is required to minimize the energy 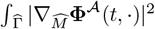, under the constraint ***v*** · ***n***_Γ_ = 0 on Γ, which is imposed using the scalar-valued Lagrange multiplier 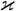. The fully discrete version of the scheme reads: given a surface 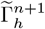 evolved from an initial surface 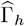, find 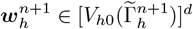 and 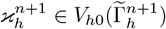 such that

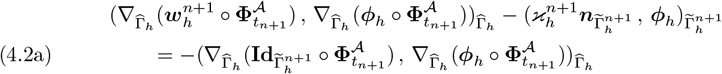

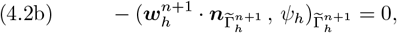

for all 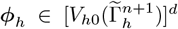 and 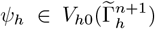. The function 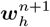 represents the displacement on 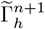 that moves tangentially the nodes maintaining good mesh properties. From this we have that 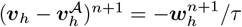.

After the surface has been displaced, a second step might be needed to extend the ALE motion in the bulk. The bulk mesh is advected through a continuous harmonic extension of 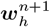. The Laplace problem reads correspondingly

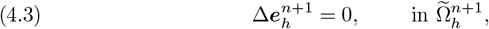

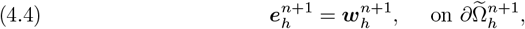

and we have that 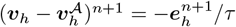.

## 5. Advection-diffusion-reaction system

The model equation we consider reads

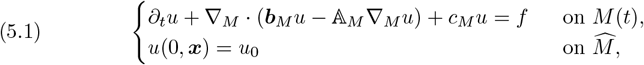

where *u* : 𝒢_*T*_ → ℝ is the concentration, ***b***_*M*_ is the advective velocity field, 𝔸_*M*_ is the diffusion matrix and *c*_*M*_ *>* 0 is the reaction constant [52, 39, 42]. As in [39, p.304] we will assume that (𝔸_*M*_)_*ij*_, (***b***_*M*_)_*i*_, ∇_*M*_ · ***b***_*M*_, *c*_*M*_ ∈ *L*^∞^(*M*), together with ***b***_*M*_ (***x***) ∈ *T*_***x***_*M, ∀****x*** ∈ *M* and the requirement for 𝔸_*M*_ to be symmetric and to map the tangent space *T*_***x***_*M* onto itself.

Extensive literature exists on finite element discretizations of such equations for what concerns the bulk case, see [46, 45]. For *closed surfaces* the SFEM discretization of parabolic PDEs in the form of (5.1) has been comprehensively reviewed in [39]. Convergence estimates for its continuous-in-time formulation using general order polynomials are found in e.g. [42]. For the sake of generality, here we focus on *moving surfaces with boundary*. The problem reads: Find *u*_*h*_ ∈ *V*_*h*_(Γ_*h*_) such that

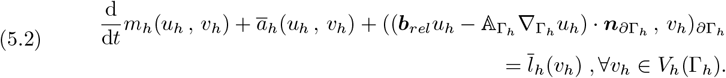

where

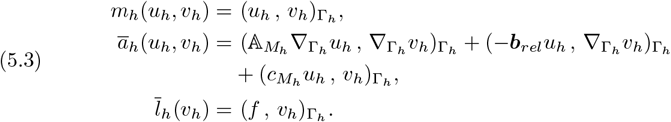

The coefficients 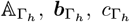 are required to satisfy the necessary smoothness conditions element-wise on the discrete surface Γ_*h*_. Recalling that 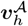 is the discrete velocity associated with the ALE mapping, the *relative velocity* is defined as 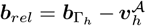. All boundary conditions are imposed weakly and the boundary is divided based on the type of boundary condition applied. We define as *∂*Γ_*h,D*_ the part of the boundary where Dirichlet conditions are applied and as *∂*Γ_*h,N*_ the part where Neumann boundary conditions are applied. It is required that

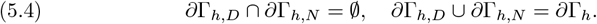

Nitsche’s boundary penalty method is used in order to impose the inhomogeneous Dirichlet boundary condition 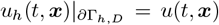weakly for the diffusive term [87, 45]. For convenience, we define *v*^+^ = max{*v*, 0} and *v*^−^ = min{*v*, 0}. The resulting bilinear forms are given by

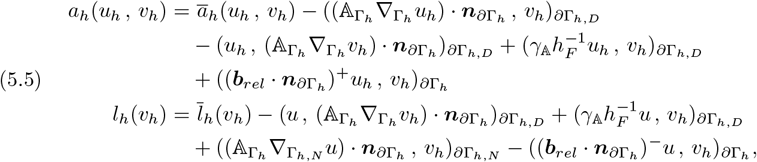

where *γ*_𝔸_ *>* 0 is the *penalty parameter* enforcing *u*_*h*_ = *u* on *∂*Γ_*h,D*_. Overall the problem for open surfaces reads: find *u*_*h*_ ∈ *V*_*h*_ such that

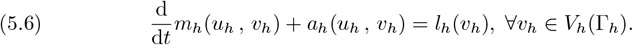

### 5.1. Numerical methods for advection-dominant problems

In biophysical cell models, as introduced in P.1, it is not uncommon to incur advection-dominant ADR equations. In our case, even purely parabolic equations might become advection-dominant due to the introduction of an ALE velocity ***v***^𝒜^. In the present work we employ and extend the continuous interior penalty (CIP) stabilization to cope with the possibly dominant advection regime. The CIP method was proposed and analyzed in [21, 20] and then later extended to the case of stationary surfaces in [23]. Moreover, the CutFEM approach for moving surfaces developed in [53] applies a CIP-type stabilization for advection dominant problems on moving surfaces. We chose CIP for the following reasons:

- It is a widely known, easily implementable technique whose implementation tools are usually shipped in classical finite element packages;
- Although only weakly consistent, it has proven to be very successful in time-dependent problems given its commutativity with the time derivative [47];
- Therefore, it has been shown to lead to convergent algorithms for discretizations of bulk, surface and implicitly described moving surfaces problems.

To formulate the CIP stabilization, we need to define averages and jumps of functions across edges and faces. For a piecewise discontinuous function *f* defined on a surface or bulk mesh 𝒯_*h*_(*M*_*h*_), we define its average and jump over an interior facet 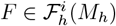 by

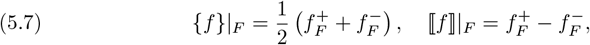

respectively, where 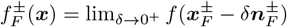.

The main idea of the CIP approach is to penalize the jump of the streamline derivative across element interfaces. To this end, the CIP stabilization form is defined as follows,

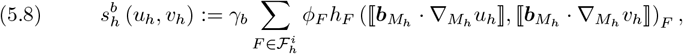

where *γ*_*b*_ *>* 0 is a dimensionless stabilization constant and *ϕ*_*F*_ denotes a stabilization parameter defined by

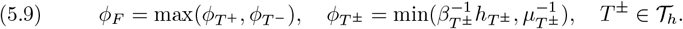

In the above *β*_*T*_ is a local velocity scale and *µ*_*T*_ is the reciprocal of a time [47]. If 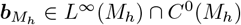, then 5.8 can be simplified to

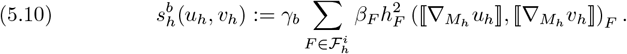

with 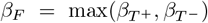. We augment the system in (5.6) further to handle advection-dominant problems, which now reads: find *u*_*h*_ ∈ *V*_*h*_ such that

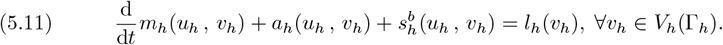

### 5.2. Numerical results for the ADR system

Given the novelty of the methods introduced, little to no theoretical convergence analysis is available in the literature and is left for future work. We thus proceed in reporting experimental convergence studies using select examples of increasing complexity to testify the accuracy of the framework.

We begin by considering a domain in 3D that moves under the linear transformation **Φ**(*t*, ***p***) = **Φ**^𝒜^ (*t*, ***p***) = 𝔸(*t*)***p*** + *B*(*t*) where

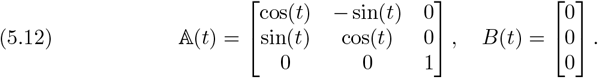

We have that **Φ**^−1^(*t*, ***x***) = [𝔸 (*t*)]^−1^(***x***−*B*(*t*)) and that the domain velocity is ***v***(*t*, ***x***) = 𝔸 (*t*)^*′*^**Φ**^−1^(*t*, ***x***) + *B*(*t*)^*′*^ ≡ ***v***^𝒜^. We construct manufactured solution for the problem

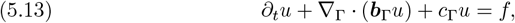

with

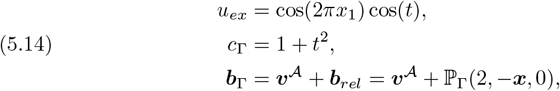

where ***x*** = (*x*_1_, *x*_2_, *x*_3_). The right-hand side *f* has been chosen so to satisfy the imposed solution. For *u*_*ex*_ we have that ∫_Γ_ *u*(*t*) = 0 and that *u* ∈ [−1, 1]. The geometry chosen is a half-sphere with unit radius that was cut along the *x*_1_-*x*_2_ plane. Under the above rotation the half-sphere has normal ***n***(*t*, ***x***) = ***x****/*∥***x***∥. We test convergence properties for the following solvers:

1. **adrSolver 1**: CIP stabilized solver 5.11.
2. **adrSolver 2**: CIP stabilized solver with imposed bound preservation in the interval [−1, 1].
3. **adrSolver 3**: CIP stabilized solver with imposed mass preservation using the lumped-mass inner product (·,·)^*h*^.
4. **adrSolver 4**: CIP stabilized solver with both imposed bound preservation in the interval [−1, 1] and mass preservation.

All solvers reveal the expected convergence in time and space. The results for **adrSolver 4** with BDF-1 time stepping are shown in Figure 3. Analogous results are obtained for the other solvers.

**Fig. 3:**
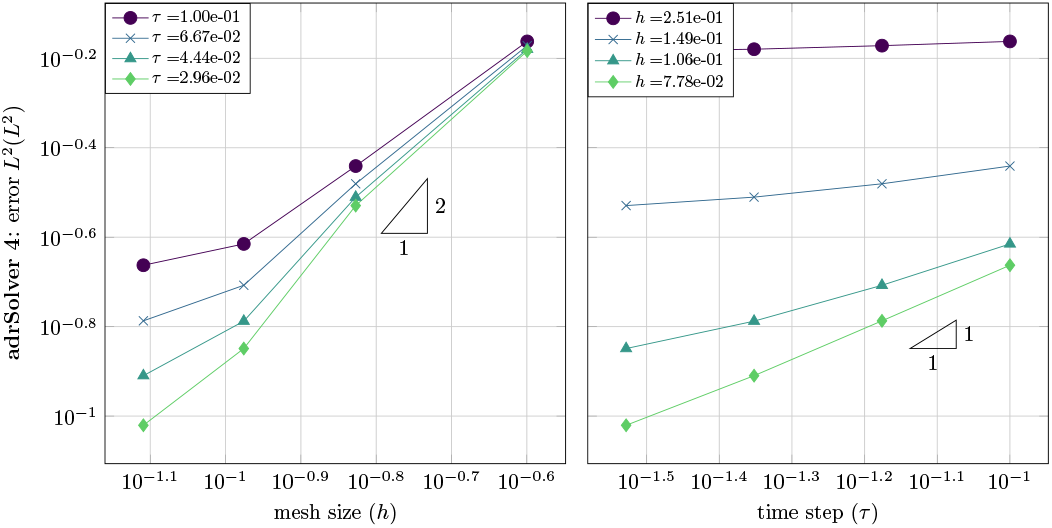
Convergence studies for the solver **adrSolver 4** in List 5.2. As expected, first and second order convergence are achieved in time and space, respectively.

To further test the effectiveness of the algorithm we now consider an ill-posed problem. Maintaining the half-sphere geometry and the transformation as in 5.12, we solve the pure transport problem

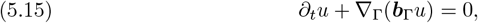

with

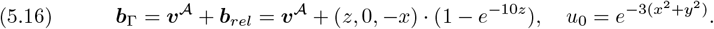

By construction the initial concentration is pushed towards the boundary where an incredibly sharp boundary layer forms, since zero flux boundary conditions are enforced by the fact that ***b***_*rel*_ ***n***_*∂*Γ_ = 0. Given the zero flux condition, the resulting solution is also mass-preserving. Parameters are set as follows
where *nv* is the number of vertices and *ne* is the number of elements. We compare the results of the four different solvers of List 5.2 in Figure 4 and Figure 5. The bounds for **adrSolver 2** and **adrSolver 4** were set to [0, 10^5^] in order to guarantee positivity.

**Fig. 4:**
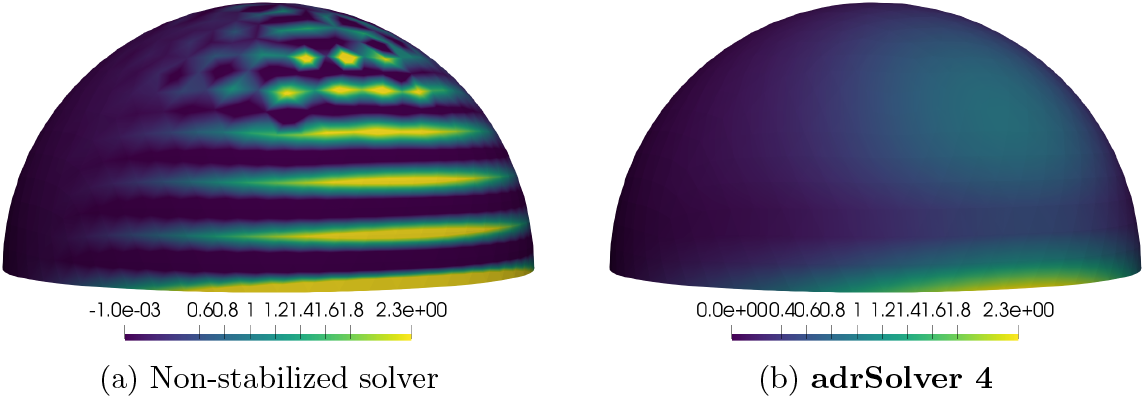
(a) Oscillating final solution of non-stabilized ADR solver at *t* = 1. (b) Final solution for **adrSolver 4** at *t* = 1.

**Fig. 5:**
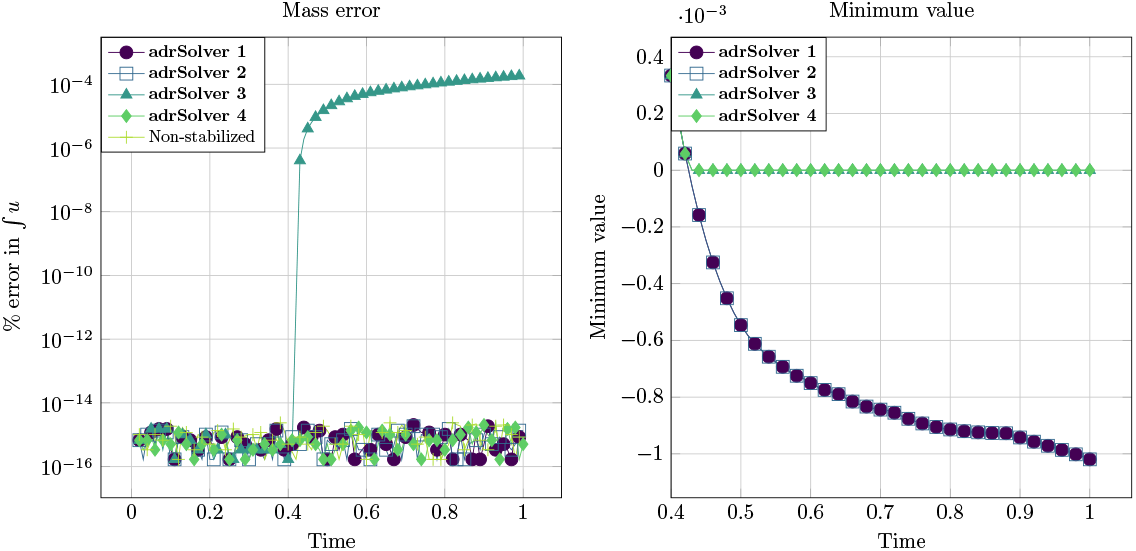
Left: relative absolute error in ∫_Γ(*t*)_ *u*(*t*) with respect to ∫_Γ(*t*)_ *u*(0) for the solvers in List 5.2. Right: Close-up of minimum value of *u* for the different solvers in List 5.2.

| $h$ | $nv$ | $ne$ | $T$ | $\tau$ |
| --- | --- | --- | --- | --- |
| 0.1 | 780 | 1495 | 1 | 0.01 |

It can be seen that the non-stabilized solver is unstable due to the pure advective regime, leading to oscillations that inevitably effect the whole domain Figure 4a. **adrSolver 1** effectively controls the solution’s gradient thanks to the stabilization, generating a non-oscillating solution, but fails to maintain its positivity, see Figure 5 (right plot). This characteristic is lost along *∂*Γ where the species encounters the steep boundary layer. **adrSolver 2** manages, thanks to the cutoff, to keep the solution positive but fails in maintaining mass-preservation as it can be seen in Figure 5 (left plot), where the relative absolute percentage mass error is plotted. **adrSolver 4** successfully manages to deal with the boundary layer, maintain positivity, and avoid oscillations. The final solution is shown in Figure 4b.

## 6. Cahn-Hilliard system as a phase separation model

Phase field modeling plays a central role in biophysical systems to investigate phase transition processes. Common phase field models for stationary surfaces are derived from Allen-Cahn and Cahn-Hilliard dynamics [26, 44, 43, 35], where the motion follows the minimization of an energy functional. We will focus here on the functional

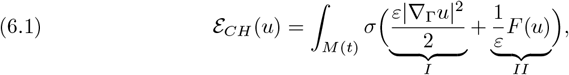

where *u* is the phase field, *ε* ∈ ℝ^+^ is a length scale parameter determining the interface length and *σ* ∈ ℝ^+^ describes the interface tension between different phases. Term *I* encodes the interface energy, penalizing inhomogeneities. Term *II* encodes the local free energy and drives phase separation through the use of a double-well potential. We consider here two different double-well potentials

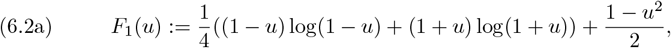

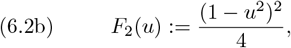

for which

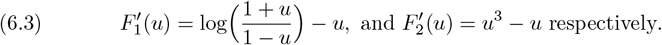

*F*_1_ is a Flory-Huggins-type mixing free energy density whose formulation finds its roots in configurational entropy [59, 48]. It can be seen that *F*_1_ is only well posed for *u* ∈ (−1, 1). For this reason, a simpler polynomial potential *F*_2_ is often used. The advantage of the latter is that it is well defined for values of *u* outside the interval (−1, 1).

The extension of the Cahn-Hilliard equation to *evolving surfaces* has seen recent developments due to its applicability to biological membranes and alloys [105, 41].

The evolving surface Cahn-Hillard equation [24, 25] can be written as

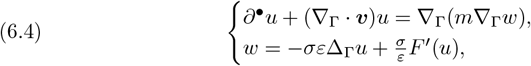

subject to the initial condition *u*(0) = *u*_0_ for suitable initial data. *w* represents the chemical potential and *m* ∈ ℝ^+^ is the mobility. As explained in [77], models for evolving surfaces have similar energy functions to models on stationary surfaces, but important differences arise. Even if the evolution of the systems is driven by the minimization of (6.1), energy dissipation is no longer satisfied since the velocity is viewed as external force that adds to the system. Analogously, mass preservation depends on the velocity. To keep mass conservation, we must assume ∇_Γ_ · ***v*** = 0, which is equivalent to assume inextensibility of the cell membrane, and by consequence area preservation. This assumption also allows to consistently extend this one-phase model to *N* -phase models satisfying the hyperlink condition, as described in [77].

### 6.1. Numerical methods for Cahn-Hilliard system

For the discretization of (6.4), the evolutionary surface finite element method (ESFEM) is one of the most popular methods adopted and the one we consider here [39]. Its advantage lies in simple implementation, low memory consumption, ability to handle complex deformations and potential for efficient parallelization. We refer to [77] for a review of alternative methods. We employ a semi-implicit approach where the fully discrete formulation reads: given 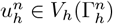 find 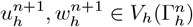 such that

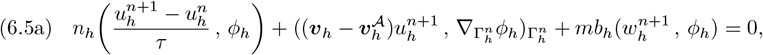

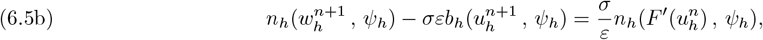

for all 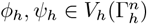 with

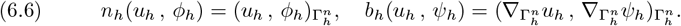

Similar discretizations can be found in [43, 7, 85], where a Cahn-Hilliard solver is coupled with a domain evolution which is unknown. In those works, the potential *F*_2_ was used. In our case, we can take advantage of the scheme presented in Section 3 to employ both *F*_1_ and *F*_2_, and apply mass conservation if needed. The scheme proceeds as follows.

In the *predictor* step we solve the unconstrained problem 6.5. Taking the non-linear term explicitly ensures 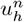 lays inside the desired bound.

In the *corrector* step we solve the constrained problem imposing 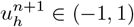.

#### Remark 6.1.

Another strength of the postprocessing technique of [29] evident here is that it can act directly on the quantity of interest *per se* without involving 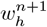, which is unbounded. We remind the reader that extensions for higher order BDFs methods are presented in the original articles.

### 6.2. Numerical results for Cahn-Hilliard system

We proceed in reporting experimental studies to assess the accuracy of the proposed framework. Consider a cylinder revolving around the *z*-axis with radius *R* = 1 and height *l* = 2. To satisfy the assumption ∇_Γ_ · ***v*** = 0, we transform the initial cylinder through the isometry **Φ**(*t*, ***p***) = **Φ**^𝒜^(*t*, ***p***) = 𝔸 (*t*)***p*** + *B*(*t*)

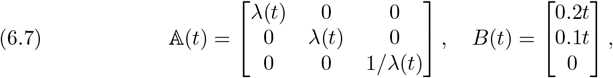

where *λ*(*t*) = 1 + 0.5 sin(*πt*). Convergence studies have already been proposed in the literature and we proceed by testing the quality of the structure preservation technique on realistic initial conditions. The verification is performed for the following solvers:

1. **chSolver 1**: Solver 6.5 with no postprocessing.
2. **chSolver 2**: Solver 6.5 with imposed bound preservation in the interval (−1, 1).
3. **chSolver 3**: Solver 6.5 with imposed mass preservation using the lumped-mass inner product (·, ·)^*h*^.
4. **chSolver 4**: Solver 6.5 with imposed bound preservation in the interval (−1, 1) and mass preservation.

Zero flux Neumann boundary conditions are imposed for both *u* and *w*. A random uniform initial distribution in the interval [−1, 1] is chosen for *u*_0_. The parameters are set as

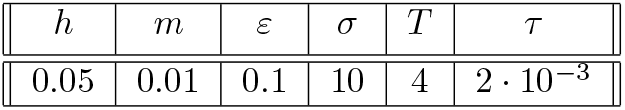

Results for the potential *F*_1_ are shown in Figure 6 and the same tests for *F*_2_ are shown in Figure 7. In Figure 6, **chSolver 1, 3** and **5** break after few steps due to the phase exceeding the bound (−1, 1), while the other solvers correctly reach the end of the simulation. Structure preservation properties are better appreciated in Figure 7. It can be seen that the energy landscape is not significantly modified by the use of either bound preservation and/or mass preservation. In both mass plots it is evident how the mass preservation property is lost for **chSolver 2**, that only has bound preservation, leading to a different overall dynamics for **chSolver 2**. Mass in instead correctly preserved by **chSolver 4** as expected. Plotting the maximum and the minimum value in the bottom two pictures not only highlights the bound preservation but also visually testify that the phase separation dynamics is respected by the postprocessing step. This is particularly clear in the passages where the bound preservation is deactivated. For example around *t* ≈ 1 in Figure 7, when the bound preservation is not needed, the maximum value of the bounded solvers naturally returns to align with the unbounded ones.

**Fig. 6:**
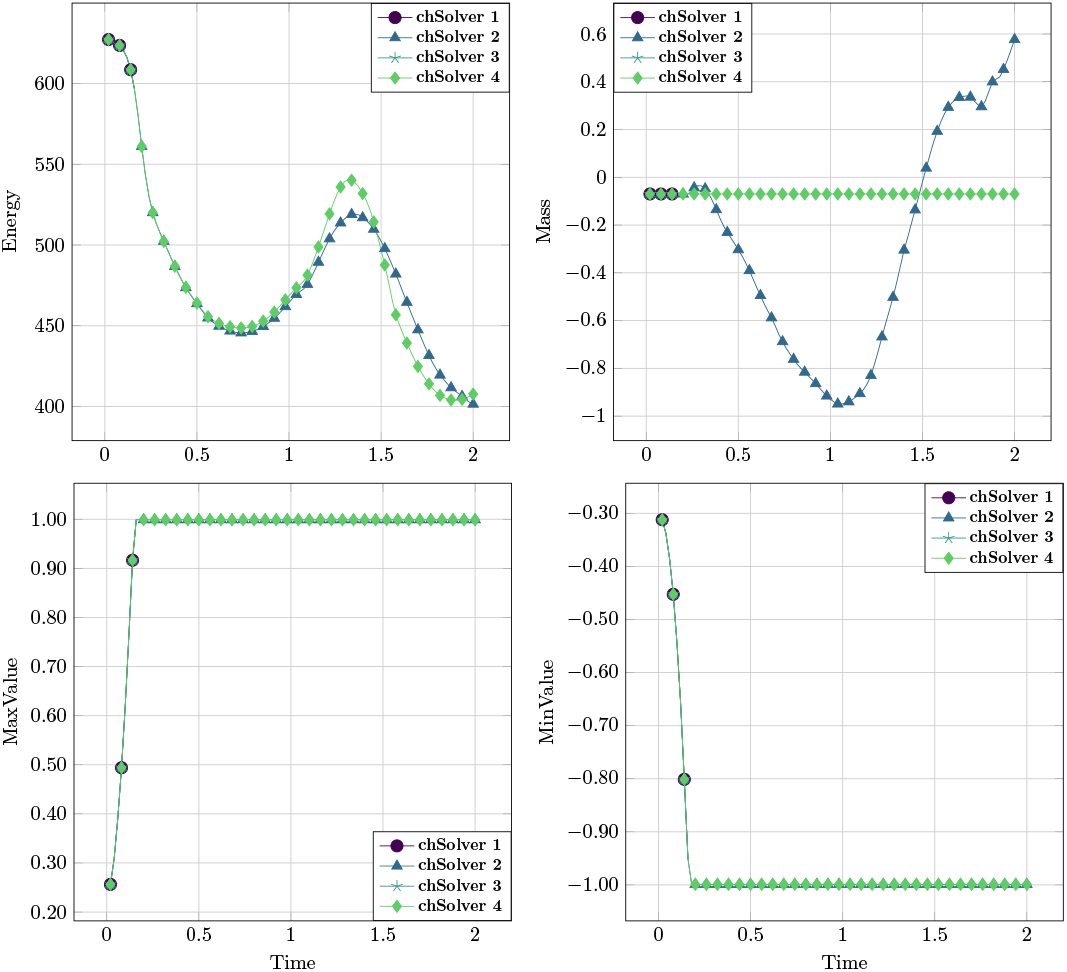
Experimental studies for potential 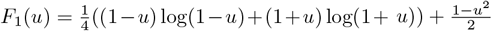and the different solvers in List 6.2.

**Fig. 7:**
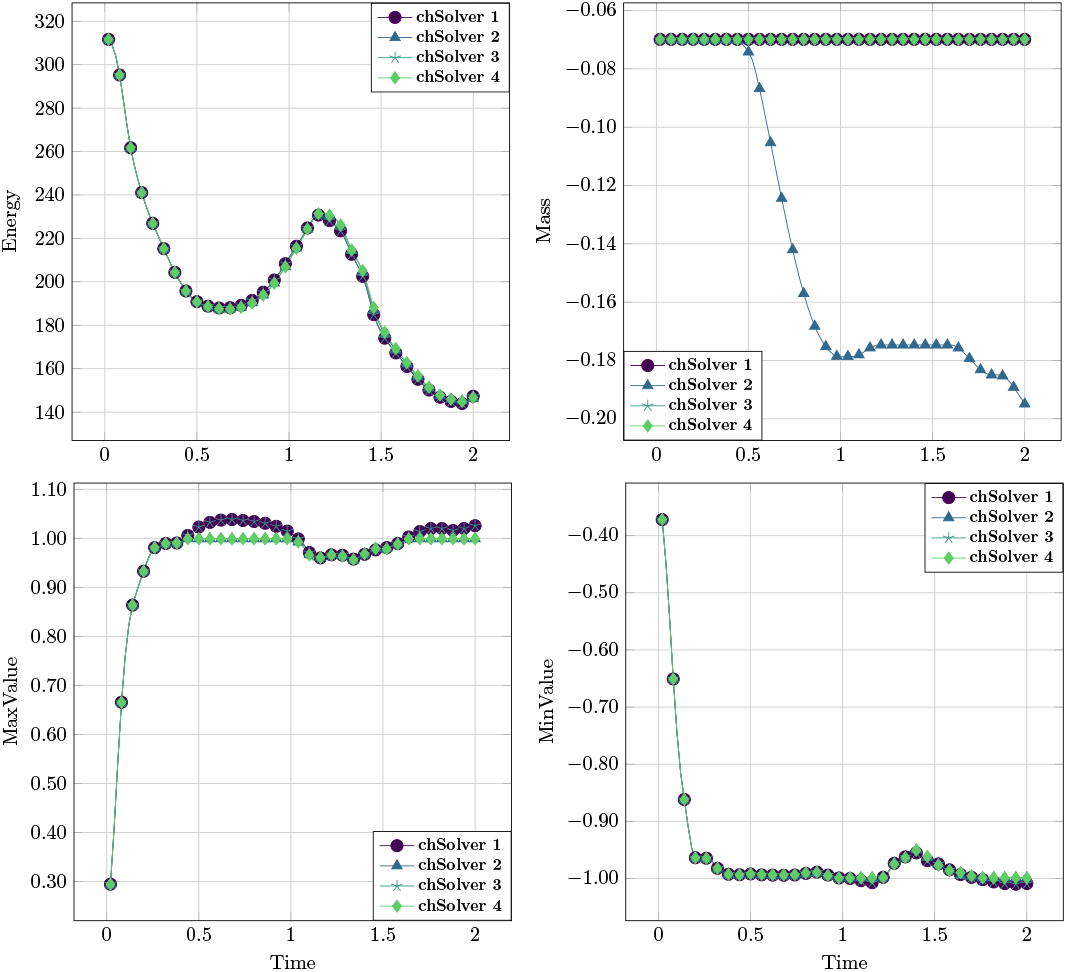
Experimental studies for potential *F*_2_(*u*) = (1− *u*^2^)^2^*/*4 and the different solvers in List 6.2.1

## 7. Membrane force system

In the realm of mathematical modeling, principal curvatures have long been used for the theory of elastic plates and shells, with the work of Kirchhoff [66] representing what may be the most famous case. Later in the ‘70s, the work of Canham [27] and Helfrich [54] on the link between biological membranes shape and curvature functionals has *de facto* initiated an entirely new field of research focused on the use of geometry of hypersurfaces for biophysical modeling.

We will here focus on the Canham-Helfrich functional *ℰ*_*B*_(*t*) and its discretization. Considering membranes with boundaries, a commonly used expression for the energy functional is

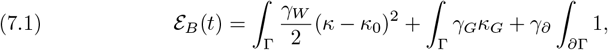

where *γ*_*W*_, *γ*_*G*_ ∈ ℝ are bending rigidities, *κ* : 𝒢_*T*_ → ℝ is the previously introduced mean curvature, *κ*_0_ ∈ ℝ is the spontaneous curvature, and *κ*_*G*_ : 𝒢_*T*_ → ℝ is the Gaussian curvature *κ*_*G*_ = det 𝕎. The parameter *γ*_∂_ takes into account the possible line energy of ∂Γ. For the sake of simplicity we will consider the case *γ*_*G*_ = *constant, γ*_*W*_ = *constant*, and assume that no topological change occur along the evolution. Taken into account these simplifications, the gradient flow of a membrane Γ is a family of evolving surfaces with normal velocity ***v***^⊥^ given by [88, 14]

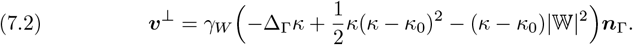

Note that such velocity is in the normal direction since tangential changes to the surface shape do not modify its energy. Equation (7.2) can be considered as a generalized Willmore flow equation. The mean curvature *κ* is directly coupled to the surface Γ through a second-order dependency. This can easily be seen in the case of surfaces whose configuration is described by a levelset equation. In that case, *κ* is the trace of the extended Weingarten map ℍ which in turn can be computed as the Hessian of the levelset for Γ. Equation (7.2) thus leads to a fourth-order nonlinear equation posed on a moving manifold.

### Remark 7.1.

Other geometrical flows like the mean curvature flow and the surface diffusion flow are also of interest in this field. Being somewhat simpler, extensive research is already available for them, while less is known about the Willmore flow, which we focus on in this article. We refer to [38, 32, 15] for further details.

### Remark 7.2.

Since this is a defining equation for the normal *velocity*, additional forces can be taken into account by introducing a right-hand side to (7.2).

### 7.1. Numerical methods for Helfrich system

We choose here the family of BGN-type algorithms as pioneered in [11] and later adapted for Willmore flow in [12, 14]. The schemes pivot around the two following ingredients:

- Introduce the *mean curvature vector* ***κ***(*t*) = *κ*(*t*)***n***_Γ_(*t*) as independent variable and use the defining equation for the mean curvature vector ***κ*** = Δ_Γ_𝕀_Γ_ in order to reduce the fourth order problem to a coupled second order one [11]; Use the weak formulation to rewrite and simplify the term *κ* |𝕎 |^2^ (more precisely *κ* |ℍ| ^2^ in the immersed setting) taking advantage of the newly introduced variable ***κ***. This reformulation not only simplifies and linearizes the system but also leads to stable schemes as proven in [37].

Our scheme choice builds on the one derived in [14] for boundary value problems. Among the different types of boundary conditions considered in [14] we will restrict to the case of *clamped boundary conditions*, for which

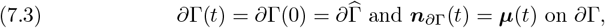

where ***µ***(*t*) is a user-given function that dictates the evolution of ***n***_∂Γ_(*t*). The weak form of (7.2) then reads: Find ***v***^⊥^, ***y, κ*** ∈ [*H*(*t*)]^*d*^ such that

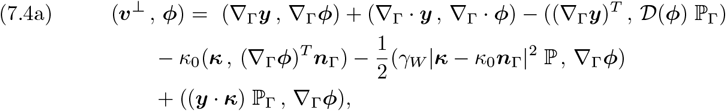

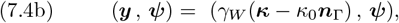

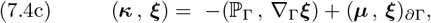

for all ***ϕ, ψ, ξ*** ∈ [*H*(*t*)]^*d*^, where [*H*(*t*)]^*d*^ is the Sobolev space *W* ^1,2^(Γ(*t*)). In the above *𝒟* (***ϕ***) = ℙ _Γ_(∇_Γ_***ϕ*** + ∇_Γ_***ϕ***^*T*^) ℙ _Γ_ and ***y*** is an auxiliary variable introduced to deal with the spontaneous curvature. For the discretization of the normal velocity in (7.4) we choose to introduce the displacement variable 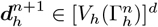 defined as

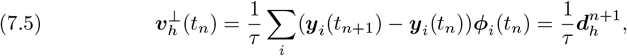

where we recall that ***y***_*i*_ are the mesh points and {***ϕ***_*i*_} the associated vector-valued basis such that ***ϕ***_*i*_(***y***_*j*_) = δ_*ij*_ at all times. Moreover, (7.4c) and (7.4b) can be collapsed

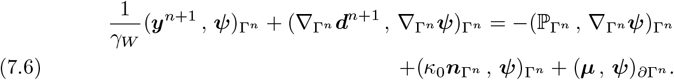

The fully discrete form of (7.4) reads: Given 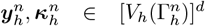 find 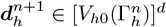 and 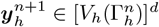 such that

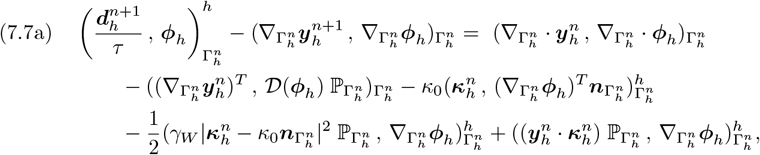

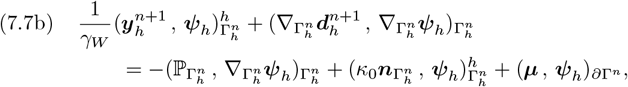

for all 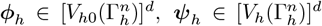, where (·, ·)^*h*^ is the mass-lumped inner product.

#### Remark 7.3.

The mean curvature 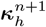 can be recovered a posteriori as 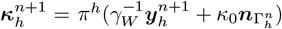 where *π*^*h*^(·) is the standard interpolation operator.

#### Remark 7.4.

Note that (7.7) corresponds to the scheme presented in [12] for the choice of the parameter *θ* = 1, i.e. without tangential motion control.

### 7.2. Numerical results for Helfrich system

To test the accuracy of our method for gradient flows, we start by performing classical studies on Willmore flow such as the ones in [37, 12, 14]. For every test we compare

1. **hfSolver 1**: algorithm (7.7).
2. **hfSolver 2**: algorithm (7.7) together with the mesh redistribution described in (4.2). We highlight that it is of crucial importance that, when the algorithm in Section 4 is employed, geometrical quantities characteristic of the mesh are updated. In our case, we refer to the mean curvature ***κ***_*h*_ and the auxiliary variable ***y***_*h*_ of the algorithm in (7.7).

We start by testing the convergence using the example presented in [12, Section 5.3]. It is an evolving sphere with initial radius *R*_0_ and spontaneous curvature *κ*_0_. The radius evolution *R*(*t*) is the solution to the ordinary differential equation

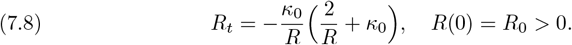

The error norm chosen to verify convergence is the following

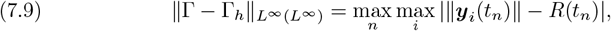

where ***y***_*i*_ are the vertices of the mesh simplices. Results are presented in Figure 8. The convergence in space and time is clearly visible and in accordance with the optimal expected. Moreover, we can see that the use of the mesh redistribution actually improves the convergence rates. We believe this behavior is due to the automatic mesh adaptivity of Equation 4.2 (already mentioned in the original article [36]), which tends to accumulate nodes in certain regions providing better geometry description.

**Fig. 8:**
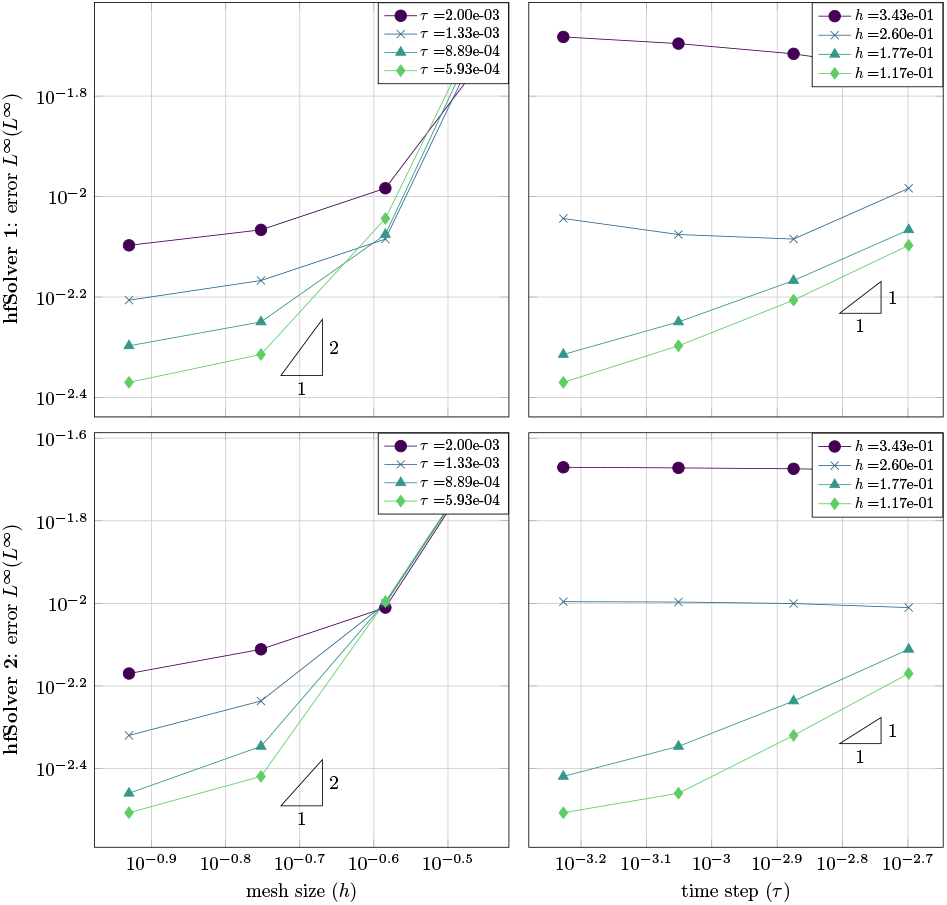
Convergence studies for Willmore flow of sphere under spontaneous mean curvature for the solvers in List 7.2.

As a second test we check the energy evolution for a torus of major radius *R* = 2 and minor radius *r* = 1 as done in [12, Section 5.1]. The minimizer of the Willmore energy in this case is the Clifford torus with radius 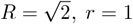, which has energy *ℰ* = 4*π*^2^ ≈ 39.4784. We use both solvers and plot the energy history for two different examples described by the following parameter sets:

| $h$ | $nv$ | $ne$ | $\tau$ | $T$ | $h$ | $nv$ | $ne$ | $\tau$ | $T$ |
| --- | --- | --- | --- | --- | --- | --- | --- | --- | --- |
| 0.2 | 2238 | 4476 | $10^{-3}$ | 2 | 0.1 | 8951 | 17902 | $10^{-3}$ | 2 |

The results are shown in Figure 9. We notice how both simulations using **hfSolver 1** break before reaching the end time. This is due to instabilities that occur at regions of high curvature already observed in [12, Figure 2] for *θ* = 1. Applying the redistribution doesn’t impact the energy landscape and allows to proceed with the simulation until the stable state is reached. Moreover, refining the mesh shows how the energy converges toward the expected value.

**Fig. 9:**
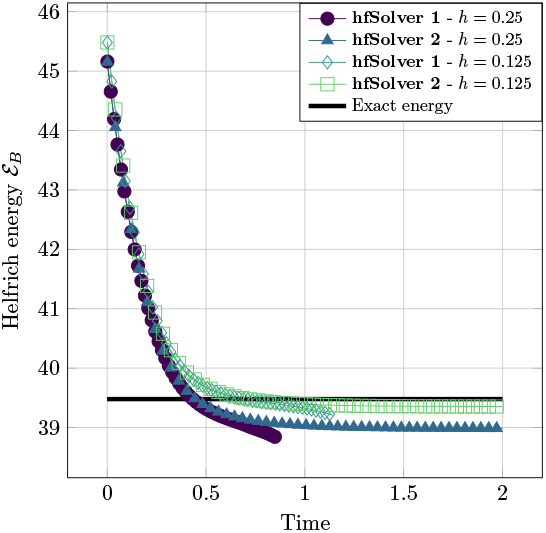
Energy evolution for torus with *R* = 2, *r* = 1 for the solvers in List 7.2.

#### 7.2.1. Pinching tests for Helfrich system

We perform consolidated tests for the Willmore flow under the influence of spontaneous mean curvature as presented in [12, Section 5.3] and then repeated in [51], where a similar two-step procedure as the one adopted here is used. We consider two different tests for cigar-shaped surfaces:

1. **hf-Test 1** with initial mesh as in Figure 10a and the following sets of parameters

2. **hf-Test 2** with initial mesh as in Figure 10d and the following sets of parameters

| $h$ | $nv$ | $ne$ | $T$ | $\kappa_0$ | $h$ | $nv$ | $ne$ | $T$ | $\kappa_0$ |
| --- | --- | --- | --- | --- | --- | --- | --- | --- | --- |
| 0.25 | 601 | 1198 | 1 | -2 | 0.125 | 2321 | 4638 | 1 | -2 |

2. **hf-Test 2** with initial mesh as in Figure 10d and the following sets of parameters
| $h$ | $nv$ | $ne$ | $T$ | $\kappa_0$ | $h$ | $nv$ | $ne$ | $T$ | $\kappa_0$ |
| --- | --- | --- | --- | --- | --- | --- | --- | --- | --- |
| 0.25 | 845 | 1686 | 0.3 | -3 | 0.125 | 3282 | 6460 | 0.3 | -3 |

A common timestep of *τ* = 10^−3^ has been used for all the experiments. The idea is to test the ability of the algorithm to simulate pinching phenomena, since **hf-Test 1** develops one neck and **hf-Test 2** develops two necks. The results are plotted in Figure 10 for both **hfSolver 1** and **2**. It can be seen how the mesh adaptivity mentioned in [36] leads to a better resolution of the neck and overall better results for **hfSolver 2**. By comparison of Figure 10c with [12, Figure 12] the shape is qualitatively in accordance with what is expected. Even more, while [12, Figure 12] requires a finer mesh to accurately resolve the three pearls, the mesh adaptivity of [36] alleviates that requirement for similar resolution. Figure 10 is also in accordance with [51, Figure 5.14], where a similar algorithm is employed for a cigar-shaped surface with different aspect ratio. As it happens in that article, it has to be noted that the refinement of the necks comes to the expense of a coarser mesh at the lobes. Looking at the energy evolution for **hf-Test 2**, the curve for **hfSolver 1** and **hfSolver 2** fall in the same place for all experiments, highlighting that the tangential mesh redistribution does not impact the energy landscape of the shape dynamics. For **hf-Test 1** we have that the dynamics is stable for both examples when *h* = 0.2, with a slight difference in energy when necking occurs. Refining the mesh leads to mesh breaking for **hfSolver 1**, while **hfSolver 2** maintains its stability, asymptotically tending towards the same energy **hfSolver 1** was tending to in the coarser case. Overall, the proposed technique appears stable in various regimes and accurate in resolving the energy evolution of the geometry.

**Fig. 10:**
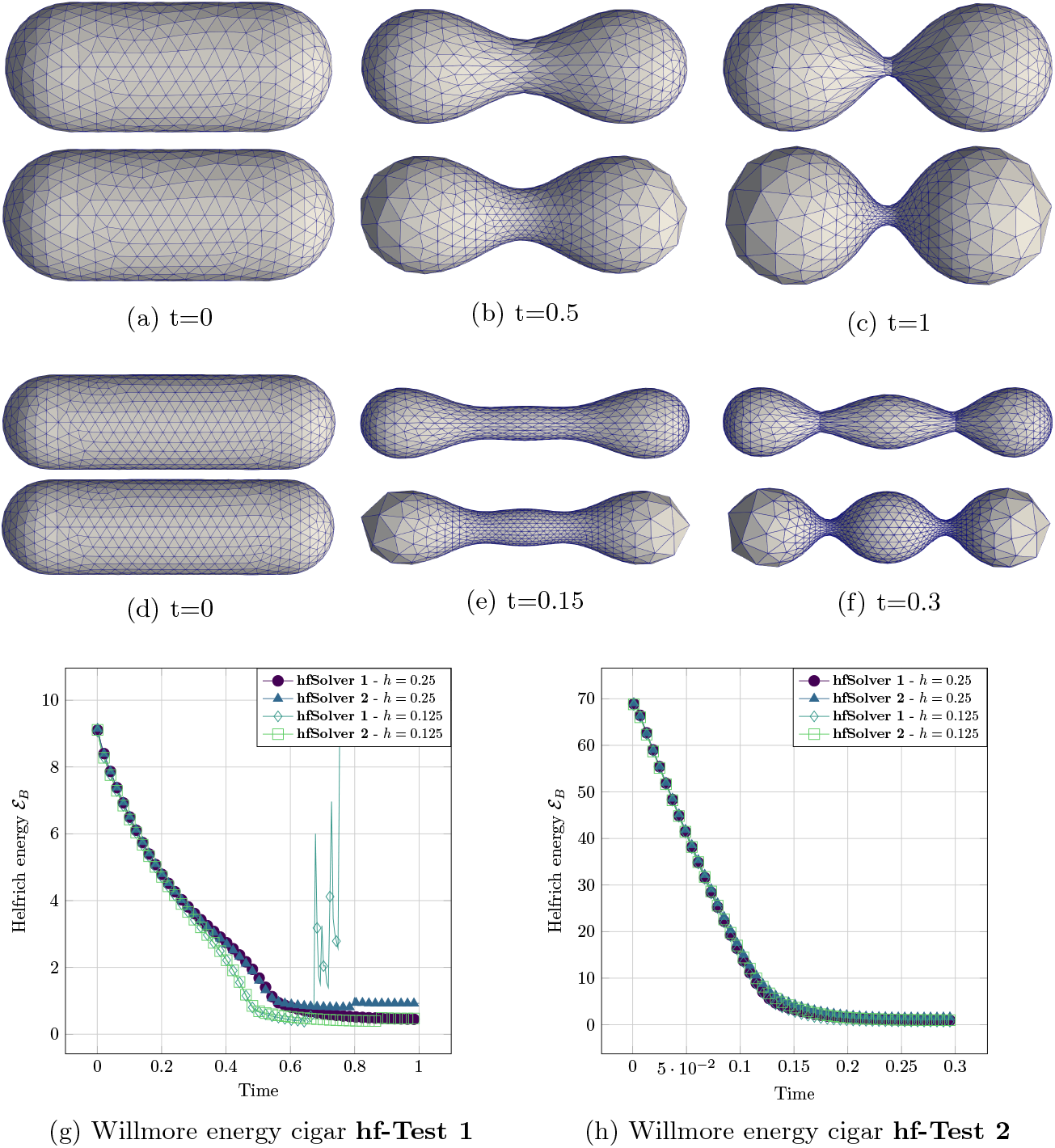
(a)-(c): Mesh evolution for **hf-Test 1** (coarser mesh). Top: **hfSolver 1**, bottom **hfSolver 2**. (d)-(f): Mesh evolution for **hf-Test 2** (coraser mesh). Top: **hfSolver 1**, bottom **hfSolver 2**. (g) Energy evolution for **hf-Test 1** and (h) Energy evolution for **hf-Test 2** for the different solvers in List 7.2. We note that figures are rescaled.

## 8. Coupling

Simulating actual biological behavior involves complex coupling between different equations. Given we make heavy use of post-processing techniques, we couple the various solvers in a staggered way. We have then the option to choose between explicit and implicit staggering. Both approaches have shown to be effective in the literature [84, 65, 64, 51]. In [7, Appendix E], where fluid deformable membranes with phase separation are considered, both procedures show very similar results even in the most complex scenarios. Moreover, when implicit staggering is chosen, the authors of [7] claim that convergence is achieved in four steps on average. Strong of these results we proceed in coupling our solvers.

### Remark 8.1.

An alternative option is to use a monolithic approach and couple the solvers in a unique system, avoiding staggering. In this way there is no need for sub-iterations and memory allocation is reduced. On the other side, the solution is usually only available for ad-hoc problems and not flexible. Together with this, eventual non-linearities have to be linearized. One can find an example of such a choice in [85], where wetting dynamics of liquid droplets on deformable membranes is simulated.

### 8.1. Numerical results for coupled mean curvature and ADR equations

We begin by testing the convergence properties of our coupled solver. To do so we pick the coupling example presented in [67]. The system which is solved is the following [67, p.686]:

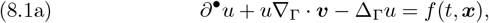

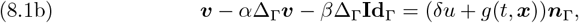

with parameters *α, β, δ* ∈ ∈ ℝ^+^ and ***x*** = (*x*_1_, *x*_2_, *x*_3_). Convergence studies are perfomed choosing *f, g* such that the exact solution for *u* is *u*(*t*, ***x***) = *x*_1_*x*_2_*e*^−6*t*^. The geometry is chosen to be a sphere whose radius evolves following the law

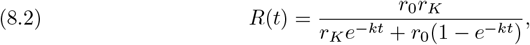

with parameters *r*_0_, *r*_*K*_, *k* ∈ ℝ^+^. The parameters are chosen as follows

|  |  |  |  |  |  |  |
| --- | --- | --- | --- | --- | --- | --- |
| $T$ | $\alpha$ | $\beta$ | $\delta$ | $r_0$ | $r_K$ | $k$ |
| 1 | 0 | 1 | 0.4 | 1 | 2 | 0.5 |

To simulate (8.1a) we can use what has been introduced in Section 5 and add a right-had side coupling. Since we chose *α* = 0, we can simulate (8.1b) by simplifying (7.7). The result is a weighted mean curvature solver that, in this very case, takes the form

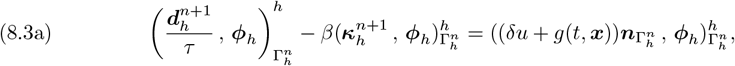

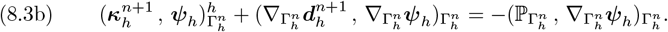

We perform the test using implicit staggering. The results of the convergence studies are shown in Figure 11, where the same norm as in Equation 7.9 is used to study the results. One can see how the expected convergence is achieved in both space and time. To qualitatively visualize the effect of the mesh redistribution and ADR stabilization, the example in [67, Sec. 11.2] is also reproduced. The simulation is derived from a proposed model for tumor growth. The coupled system is as follows:

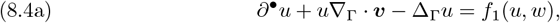

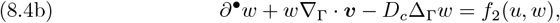

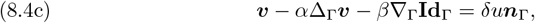

with the non-linear couplings

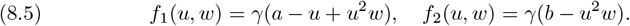

**Fig. 11:**
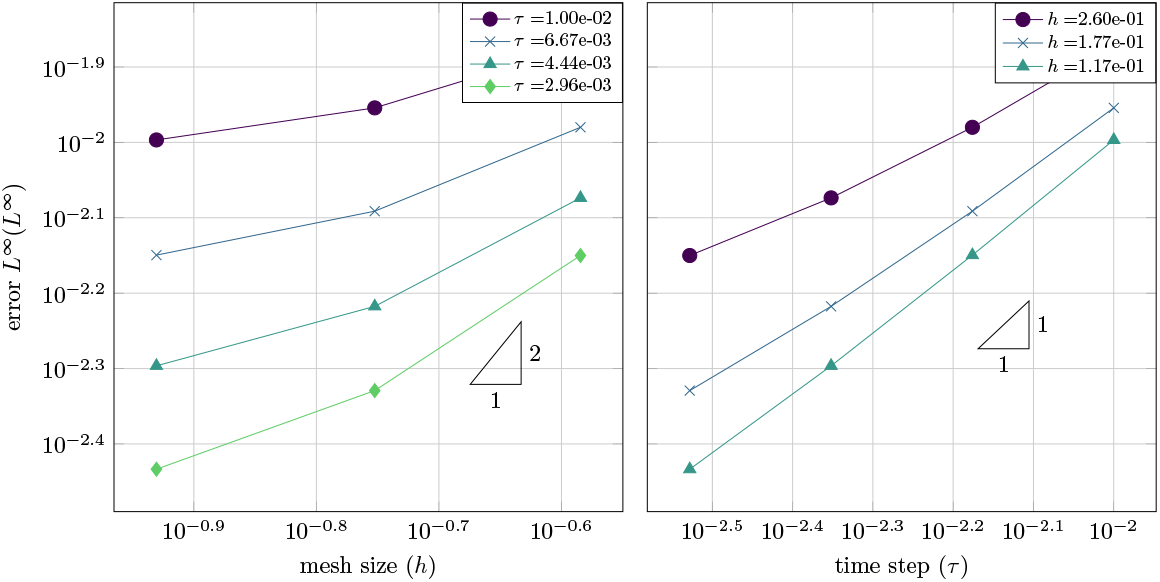
Convergence studies for **mcadrSolver 4** and problem (8.1).

The parameters are set as

| $T$ | $\tau$ | $h$ | $\alpha$ | $\beta$ | $\delta$ | a | b | $\gamma$ | $D_c$ |
| --- | --- | --- | --- | --- | --- | --- | --- | --- | --- |
| 8 | $10^{-3}$ | 0.07 | 0 | 0.01 | 0.4 | 0.1 | 0.9 | 100 | 10 |

As discussed in [67], the nonlinear system composed by (8.4a) and (8.4b) is solved until *t* = 5 without coupling it to the mesh evolution, i.e. ignoring (8.4c). After that, the full system is evolved until *t* = 8. Implicit staggering is used. We thus have the following solvers:

1. **mcadrSolver 1**: Implicit staggering of (5.6) and (8.3) and no mesh redistribution.
2. **mcadrSolver 2**: Implicit staggering of (5.11) and (8.3) and mesh redistribution.

In Figure 12 we compare the mesh quality at the final time *t* = 8. It is evident that the redistribution step keeps the mesh well-behaved while maintaining solution accuracy.

**Fig. 12:**
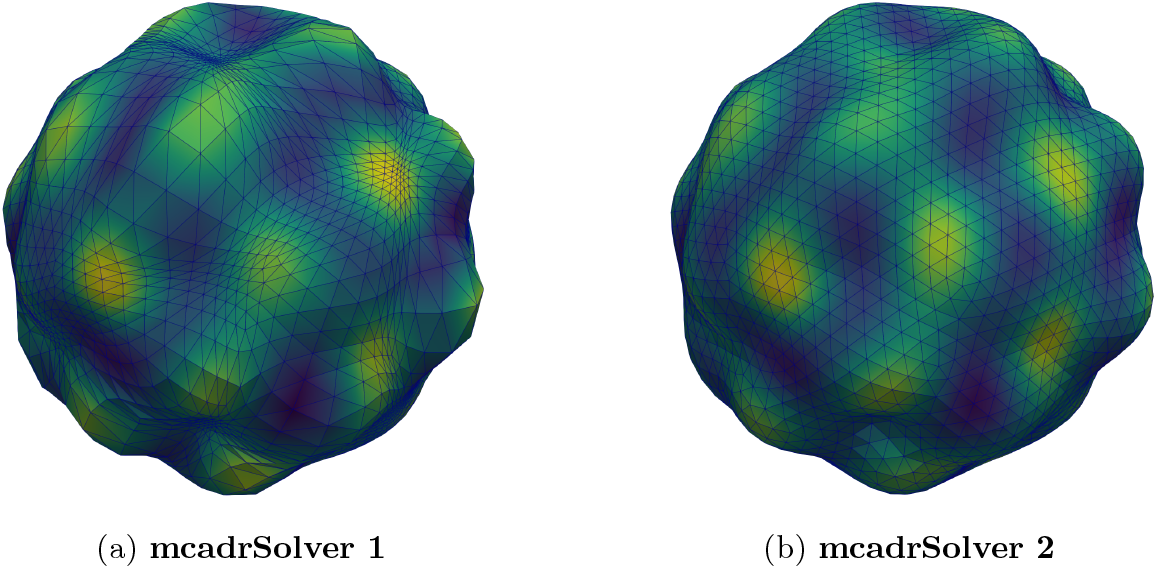
Simulation at *t* = 8 for the model (8.4) [67, Sec. 11.2] and the implicitly staggered schemes in List 8.1.

### 8.2. Numerical results for coupled Helfrich flow and Cahn-Hilliard euqations

We proceed now to test the coupling for more complicated systems and geometry evolution to explore the potential of our framework. The overdamped limit model for fluid deformable two-component membranes presented in [7, p.A41-33] is considered. The discrete scheme (7.7) is augmented with a Lagrange multiplier to maintain the inextensibility constraint 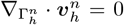. The modified version reads: Given 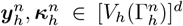 find 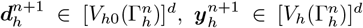 and 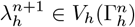 such that

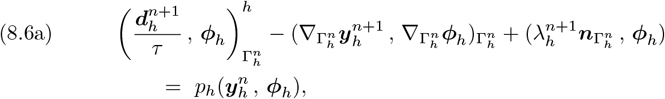

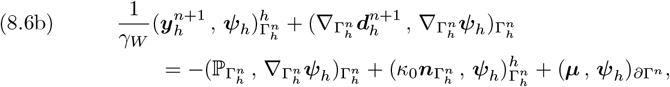

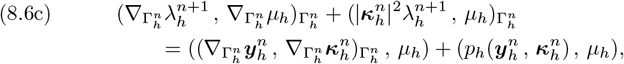

for all 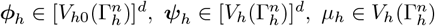 where

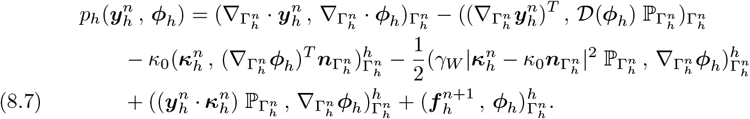

In (8.7), 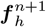 is a general right-hand side. The system is composed of the following three steps:

1. Solver (8.6) with the right-hand side (recall (6.1)):

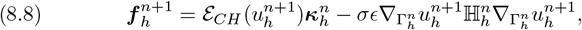

where 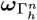 is a certain vertex normal, see [11] for details, and 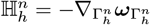 is the discrete extended Weingarten map.
2. Solver (4.2) where the prescribed material velocity is set to

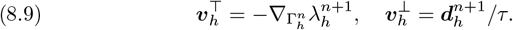
3. Solver (6.5) with tangential advective velocity 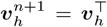 and polynomial potential *F*_2_ (6.2).

The test considers a sphere of radius one and the following initial parameters:

| $T$ | $h$ | $\sigma$ | $\varepsilon$ | $m$ |
| --- | --- | --- | --- | --- |
| 1 | 0.08 | $1.5\sqrt{2}$ | 0.1 | 0.001 |

The common initial condition for the Cahn-Hilliard solvers is

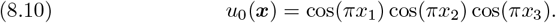

The simulation is run with variable elasticity modulus *γ*_*W*_ ∈ {0.5, 0.1, 0.02} following the parameters in [7]. The timestep is varied accordingly with values *τ* ∈ {10^−4^, 10^−4^, 2.5 ·10^−5^} respectively. The simulation is stopped once the area difference between initial and evolved surface differs by more than 10%. Results are presented in Figure 13, where it can be seen that the characteristic bulging dynamics is correctly recovered.

**Fig. 13:**
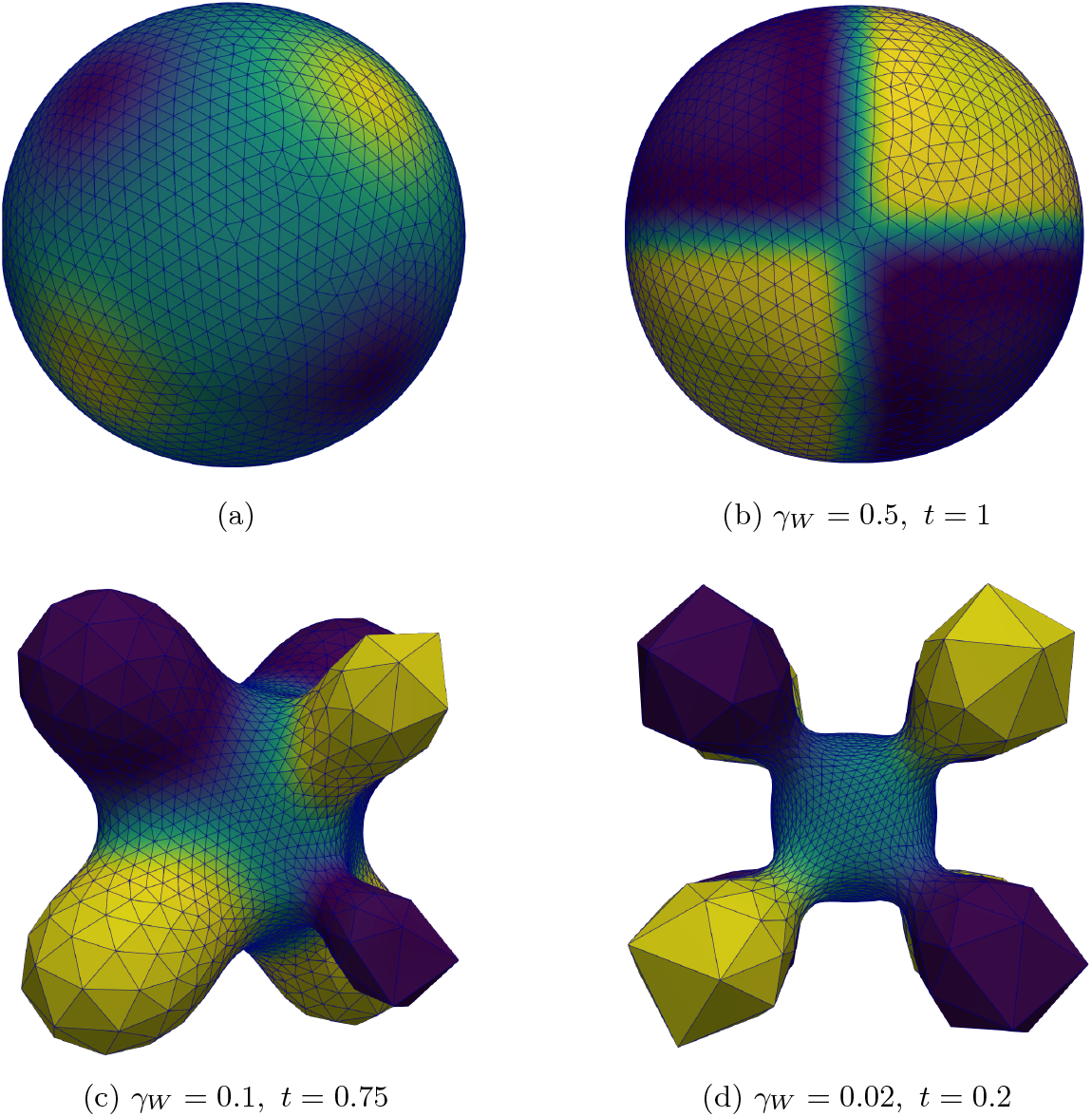
(a) Initial setup for problem the problem 8.10. (b)-(d) Mesh for different elasticity modulus. The simulation is shown at end time *t* = 1 or at last timestep before inextensibility threshold is violated.

## 9. Conclusions and outlook

Here, we presented a numerical framework to simulate moving boundary problems in biophysics. This work addresses a longstanding need for robust computational frameworks to tackle such problems. While other frameworks exist [85, 6, 7, 83, 13], our work targets flexibility of application while maintaining accuracy. Algorithms that enjoy stability and convergence properties are tied to non-invasive postprocessing techniques so to favor complex applications. The result is a stable pipeline with the capability of being scaled to more realistic scenarios.

Specifically, structure preservation is introduced on moving bulk and surface domains as a mean to allow for biophysically justified models and guarantee interpretability. We show that the scheme is readily implementable while being fast and accurate. The flexibility of the algorithm, which does not require *ad hoc* implementation, is proven by applying it to both advection-dominant ADR equations and phase-field models of the Cahn-Hilliard type. Convergence studies were performed for the bound- and mass-preserving scheme and the expected convergence rates were numerically verified. The framework also couples recent mesh redistribution techniques for gradient flows with the ALE method. We showed that this union is efficient in handling complex reshaping dynamics without the need for remeshing. As a result, the algorithm is tunable to the needs of the user and does not modify the underlying domain evolution. This is crucial in biophysical settings where the dynamics of the overall system is strongly coupled to the domains’ shape. Convergence studies were performed, demonstrating the accuracy of the scheme. Additionally, complex benchmark tests were conducted, showing good agreement with previous results in the literature. Building on previous results, we show that staggering is an effective method to simulate increasingly complex physics such as those that occur in biophysics.

The framework developed here has strong potential to simulate problems with increasing biological complexity. To achieve this goal, we foresee the need to incorporate large systems of ADR equations, that are not discussed in the present work [50, 83, 82]. Furthermore, fluid equations have also been shown to play an important role at certain timescales [94, 96, 2]. Including Navier-Stokes type equations on both interfaces and bulk [85, 7] will broaden the applicability of the computational schemes presented here.

In summary, we presented a finite element framework able to handle a rich set of reshaping dynamics arising from bulk-surface coupled PDEs in biophysics. This framework poses itself as a major stepping stone on the path to physical simulations of moving boundary problems in cell biology.

## Acknowledgments

We acknowledge Dr. Emmet Francis for his valuable input on biologically relevant models for computational biology and for sharing the lessons learned from his programming experience [50].

